# A preliminary investigation of major bacterial antibiotic resistance genes present in Australian fecal samples using shotgun metagenomics and targeted sequencing

**DOI:** 10.1101/2020.03.19.999250

**Authors:** Vanina Guernier-Cambert, Anthony Chamings, Fiona Collier, Soren Alexandersen

## Abstract

The gut microbiota is an immense reservoir of antimicrobial resistance genes (ARGs); however, in Australia the profile of the gut ‘resistome’, or ensemble of ARGs, has not been investigated. This study provides a first preliminary mapping of the major bacterial ARGs present in human, domestic dog and wild duck fecal samples collected from south-eastern Victoria, Australia; and evaluates the use of shotgun metagenomics sequencing (SMS) and targeted amplification of ARGs. We analysed SMS data using an in-house method and web-based bioinformatics tools: ResFinder and KmerResistance. We examined targeted sequences using One Codex or the PanBacterialAnalysis Torrent Suite plugin. All methods detected ARGs in all samples, with resistance to up to 13 classes of antibiotics detected overall. ARGs were more abundant in the human and dog samples than the duck samples. They mostly conferred resistance to three classes of antibiotics that are the most frequently prescribed in Australia: tetracycline, β-lactams and MLS_B_ (macrolide, lincosamide, streptogramin B). Targeted sequencing significantly improved sensitivity for detection of ARGs included in the panel; however, SMS provided quantitative information and allowed tentative identification of the host bacteria. For SMS, web-based and in-house methods gave comparable results, with discrepancies mostly due to different reference databases. The in-house method allowed manually checking results and potential errors, while web-based methods were user-friendlier and less time-consuming. More samples need to be investigated to fully describe the resistome in humans and animals in Australia.

## Introduction

Antimicrobial resistance (AMR) occurs naturally in bacteria and other microorganisms, but increases when antimicrobials (antibiotics, antivirals, antifungals) are used^1^. According to WHO, AMR accounts for an estimated 700,000 deaths per year and is one of the ten threats to global health in 2019^2^. Reservoirs of multi drug-resistant (MDR) bacteria are ubiquitous, and the concept of an “antibiotic resistome” was first coined to describe the large collection of bacterial antimicrobial resistance genes (ARGs) found in a specific environment^3^. Although AMR is most clinically relevant in pathogenic bacteria, ARGs exist in both pathogenic and non-pathogenic bacteria, and large reservoirs of these genes exist in all ecosystems, including in commensals at all sites of the human or animal body^4^.

The adult human gastrointestinal tract harbours up to 1,000 molecular species or phylotypes often referred to as operational taxonomic units (OTUs) in 16S rRNA metagenomics^5,6^. This vast array of resident bacteria responds to environmental conditions inside the host, including altered dietary intake and antibiotic-induced disturbances. That can result in modifications in the microbial community structure and genetics, including the enrichment of ARGs within the gut microbiota. Recent studies have suggested potential links between the gut resistome and the spread of AMR: horizontal transfer of ARGs from commensal bacteria mediated by mobile genetic elements may affect potentially pathogenic organisms^7-9^. Although the development of culture-independent sequencing techniques has expanded our knowledge of the different reservoirs of ARGs, it is currently unclear how widespread commensal bacteria carrying ARGs are in the gut microbiota, and how they develop over time^10^.

There is however considerable evidence that antibiotic exposure promotes the development of a resistome in the gut microbiota^11-13^. Different features such as class, potency, spectrum and regimen of the antibiotics can influence the observed pattern of resistance of the gut microbiota^13^. Geographical differences also exist between countries in relation to the ARGs detected in human fecal samples^14^. For example, the number and abundance of ARGs was higher in the fecal samples of Chinese individuals than in the Spanish and Danish groups studied^15^. Variation in the total usage of antibiotics between countries may be one explaining factor for those differences. Indeed, both antibiotic exposure and the number of ARGs were higher in individuals from Spain, France and Italy than in individuals from Denmark, Japan and the United States^16^. In other countries such as Australia, very little is known about the resistome of humans and animals, with available studies focusing primarily on ARGs in pathogenic bacteria (e.g. *Campylobacter* or *Escherichia coli*) or in environmental samples, e.g. studying the soil resistome after swine, cattle or poultry manure usage^17-19^. One recent study however assessed the diversity and abundance of ARGs transcribed in the fecal microbiota of water birds of Australia^20^, but currently no study focused on the human fecal microbiota and its associated resistome in Australia. This clearly represents a significant knowledge gap. Regular surveillance of the resistome in Australia could be used to monitor trends in AMR over time and the emergence of resistance alleles. Resistome surveillance could also inform future prescribing guidelines for doctors in hospitals and community clinics, and veterinarians in companion and production animal settings, to help preserve the effectiveness of valuable antibiotics.

The declining costs of metagenomic sequencing technologies has led to increased molecular investigation of ARGs and has enabled a shift from phenotype to genotype-based approaches to investigate AMR^21,22^. The ultimate goal of ARG sequencing and related bioinformatic analyses is the accurate detection of the resistome and prediction of the antibiogram from genomic and metagenomic data^23^. Several major databases provide collections of ARGs covering broad categories of AMR mechanisms including the Comprehensive Antibiotic Resistance Database (CARD)^24^, Antibiotic Resistance Gene-ANNOTation (ARG-ANNOT)^25^, the Bacterial Antimicrobial Resistance Reference Gene Database^26^ hosted by the National Center for Biotechnology Information (NCBI) as part of the National Database of Antibiotic Resistant Organisms (NDARO), and ResFinder^27^ (see ^23^ for a more exhaustive list of available AMR bioinformatics software, databases, and data-sharing resources). These resources and their associated prediction tools enable detection and annotation of specific AMR elements in sequenced samples by comparing user-submitted sequences against representative sets of ARGs sequences. While identifying ARGs from a bacterial isolate from a sick patient is reasonably straightforward using whole genome sequencing, the exhaustive description of a resistome in complex environments with high bacteria diversity and relatively low abundances of ARGs is challenging. Two different strategies have been developed to study AMR in complex environments like the gut. Deep shotgun metagenomic sequencing is used to produce millions of reads that are filtered to map the ARGs. However, because reads corresponding to ARGs can be low (e.g. less than 2% of the total sequencing data in ^28^), targeted approaches (including an initial amplification of a panel of ARGs) with increased sensitivity have been developed. As new methods and new commercial panels become available, it is currently unclear what method is best to study resistomes when no pre-existing information is available, e.g. in Australia.

We investigated for the first time the diversity and abundance of ARGs in a broad range of fecal microbiota samples (a number of infants, healthy pregnant women, domestic dog, wild ducks) collected in south-eastern Victoria, Australia. We used both non-targeted NGS (analyzed in-house and with web-based tools) and targeted NGS (using two commercially available panels) and assessed and compared the performance of these tools for the analysis of bacterial resistomes in an Australian context. This preliminary study will pave the way for a broader analysis of bacterial resistomes in humans and animals in Australia.

## Methods

### Sample collection and processing

#### Sample collection

Human and animal samplings were performed in accordance with relevant guidelines and regulations. We collected stool samples from two infants and seven adults from the southeast of Australia (Table 1). The children were: one infant sampled at 4 weeks of age (ST5-1mo) and again at 18 months (ST5-18mo), and another infant sampled at 3 months of age (ST4-3mo). The study involving these samples was deemed negligible or low risk by the Barwon Health Human Research Ethics Committee and therefore exempt from full committee review (HREC approval 17/119). The adult samples originated from pregnant women at 36-weeks gestation between 2010 and 2013 with informed consent for study participation as part of the Barwon Infant Study (BIS), a birth cohort study–eligibility criteria and cohort profile are described elsewhere^29^. Ethics approval for this study was granted by the Barwon Health Human Research and Ethics Committee (HREC 10/24). All human stool samples were stored at −80°C prior to DNA extraction.

**Table 1.** List of tested human and animal samples. (n = 13). Fecal samples were DNA extracted using two different methods, as described.

| Sample ID | Host | Extraction method |
| --- | --- | --- |
| ST5-1mo | Human - child 1 month of age | PowerSoil® DNA Isolation Kit - Mo Bio |
| ST4-3mo | Human - child 3 months of age | QIAamp Fast DNA Stool Mini Kit |
| ST5-18mo | Human - child 18 months of age | QIAamp Fast DNA Stool Mini Kit |
| HS21 | Human - adult | PowerSoil® DNA Isolation Kit - Mo Bio |
| HS22 | Human - adult | PowerSoil® DNA Isolation Kit - Mo Bio |
| HS23 | Human - adult | PowerSoil® DNA Isolation Kit - Mo Bio |
| HS24 | Human - adult | PowerSoil® DNA Isolation Kit - Mo Bio |
| HS25 | Human - adult | PowerSoil® DNA Isolation Kit - Mo Bio |
| HS26 | Human - adult | PowerSoil® DNA Isolation Kit - Mo Bio |
| HS28 | Human - adult | PowerSoil® DNA Isolation Kit - Mo Bio |
| ST1-MAD* | Pool from 6 Pacific black ducks (*Anas superciliosa*) | QIAamp Fast DNA Stool Mini Kit |
| ST3-MUD* | Muscovy duck (*Cairina moschata*) | QIAamp Fast DNA Stool Mini Kit |
| DFS** | Pool from 2 dogs | QIAamp Fast DNA Stool Mini Kit |

HS: human sample; DFS: dog fecal sample. \* See <sup>30</sup>. \*\* See <sup>31</sup>.

Samples from ducks and dogs were collected in the south-eastern part of Victoria, Australia, as part of other projects investigating fecal microorganisms. Fresh fecal samples from one Muscovy duck (MUD) and a pooled sample from six juvenile Pacific black ducks (MAD) were collected in November and December 2016 respectively^30^. Bird sample collection was approved under Deakin University’s Animal Ethics Committee project number B43–2016 and Department of Environment, Land, Water and Planning permit number 1008206.

A pooled dog fecal sample (DFS) originating from two pups (7 weeks of age) was collected from the ground immediately after being deposited, with the consent of the owner^31^. These puppies were healthy at the time of collection, but they had been sick just 1-2 weeks before and an astrovirus was detected in the 7-week sample^31^. However, they were originally thought to have a *Campylobacter* infection and were likely treated with antibiotics at 5 weeks of age. Swabs from dog samples were placed in Universal Transport Medium (UTM) and stored on ice overnight. The following morning, swabs were aliquoted and frozen at −80°C until DNA extraction.

#### DNA extraction

DNA was extracted from fecal samples and swabs using the Qiagen Powersoil® DNA Isolation Kit (Cat#12888-100) (QIAGEN Pty Ltd., Victoria, Australia) or the QIAamp Fast DNA Stool Mini Kit with MoBio bead shearing (Mo Bio, Carlsbad, CA USA). The manufacturer’s protocol was slightly modified following optimization: for the inhibitors’ removal steps, 375 μL of Solution C2 and 335 μL of Solution C3 were used instead of the prescribed volumes. DNA concentrations were measured using a NanoDrop™ spectrophotometer (Thermo Fisher Scientific, Scoresby, VIC, Australia), and the quantity and size distribution of the fragments after DNA extraction was visualized on an Agilent Bioanalyzer DNA 7500 chip (Agilent Technologies) using the High sensitivity DNA kit (Agilent Technologies).

### Library preparation and sequencing

Extracted DNA was sequenced using three different methods relying on Next Generation Sequencing (NGS). We used a non-targeted sequencing of sheared DNA, hereafter referred to as “Shotgun Metagenomic Sequencing” (SMS) method, as well as a targeted metagenomic sequencing method using two commercially available Ion Ampliseq™ (PCR) panels (Life Technologies).

#### Non-targeted sequencing: Shotgun Metagenomic Sequencing

Genomic libraries were prepared separately for each genomic sample from 100 ng of DNA. DNA was fragmented using the Ion Shear™ Plus Reagents (Life Technologies, Grand Island, NY). Aiming for a 200 base-read library, the chosen reaction time for the enzymatic incubation at 37°C was 15 minutes, as advised in the manufacturer’s protocol. Sheared DNA was purified with Agencourt AMPure® XP Reagent (1.8x sample volume) (Beckman Coulter, Lane Cove, NSW, Australia). The quantity and size of sheared material was visualized on an Agilent Bioanalyzer DNA 7500 chip (Agilent Technologies) using the High sensitivity DNA kit (Agilent Technologies) assuming a targeted 200 base-read library.

The adapter ligation and nick repair were performed using the Ion Plus Fragment Library kit and Ion Xpress barcode adapters following the manufacturer’s recommendations (Life Technologies). Ligated and nick repaired DNA was purified with Agencourt AMPure® XP Reagent (1.4x sample volume) (Beckman Coulter) assuming a targeted 200 base-read library. The ligated and nick repaired DNA was size-selected individually with the E-Gel® SizeSelect™ Agarose Gel (Life Technologies). The size selected (and unamplified) libraries were quantified as per the manufacturer’s instructions using the Ion Library qPCR Quantitation Kit (Life Technologies) to determine if library amplification was required.

If required (i.e. final library quantity was < 50 pM), the size selected libraries were further amplified using Platinum® PCR SuperMix High Fidelity and Library Amplification Primer Mix as per manufacturer’s instructions (Life Technologies). This was necessary for four samples (ST4-3mo, MUD, MAD and DFS), which were amplified for 10 PCR cycles prior to purification and quantification.

All libraries were standardized to a concentration of 100 pM. Libraries were then pooled and diluted to 50 pM prior to loading onto Ion 530™ or 540™ Chips using the Ion Chef Instrument and Ion 530™ or Ion 540™ Kit (Thermo Fisher Scientific). Following template preparation, the chips were run on the Ion Torrent S5xl System (Thermo Fisher Scientific) as per company protocols. NGS and associated reactions were performed at the Geelong Centre for Emerging Infectious Diseases (GCEID), Geelong, Victoria, Australia.

#### Targeted sequencing: IonAmpliSeq™ panels

We used two commercially available community panels designed to target AMR determinants, both kindly donated by Life Technologies. The first panel used was the Ion AmpliSeq™ Pan-Bacterial Research panel, consisting of two primer pools: the one pool contained 269 amplicons targeting 21 specific bacterial species and 716 amplicons targeting 364 known ARGs; the other pool was comprised of 24 amplicons for 16S rRNA gene profiling of up to approximately 400,000 16S rRNA sequences. From this panel, we only used the first primer pool targeting the ARGs. The second panel used was the Ion AmpliSeq™ Antimicrobial Resistance (AMR) Research panel that also consisted of two primer pools comprising a total of 815 amplicons targeting 478 known ARGs^32^. Both primer pools were used from this panel. The list of primers is available upon request from Life Technologies (https://www.thermofisher.com/).

Ten nanograms of extracted DNA were PCR-amplified with the Ion AmpliSeq™ Library Kit (Life Technologies) and Ion AmpliSeq™ Panels, both panels including a 5X Ion AmpliSeq™ HiFi Master Mix. With the Pan-Bacterial Research panel, PCR conditions were as follows: enzyme activation at 99°C for 2 minutes followed by 18 cycles of 99°C for 15 seconds and 60°C for 8 minutes before holding at 10°C. With the AMR Research panel, PCR conditions were as follows: enzyme activation at 99°C for 2 minutes followed by 19 cycles of 99°C for 15 seconds and 60°C for 4 minutes before holding at 10°C. The following steps (adaptors and barcodes ligation, purification, and qPCR quantification) were done as per the manufacturer’s instructions. None of the AmpliSeq™ libraries required further amplification at this stage. Libraries were then pooled prior to loading onto an Ion 530™ Chip and loading into the Ion Chef Instrument. Following template preparation, the chip was run on the Ion Torrent S5xl System (Thermo Fisher Scientific) following company protocols. We submitted reads obtained from the SMS and targeted sequencing to the European Nucleotide Archive (ENA) under project accession number PRJEB36405.

### Next generation sequence analyses

We obtained three sets of sequence data that we analysed using different tools and methods. A schematic flowchart summarizing the different sequencing and analysis methods is presented in Figure 1.

**Figure 1.**
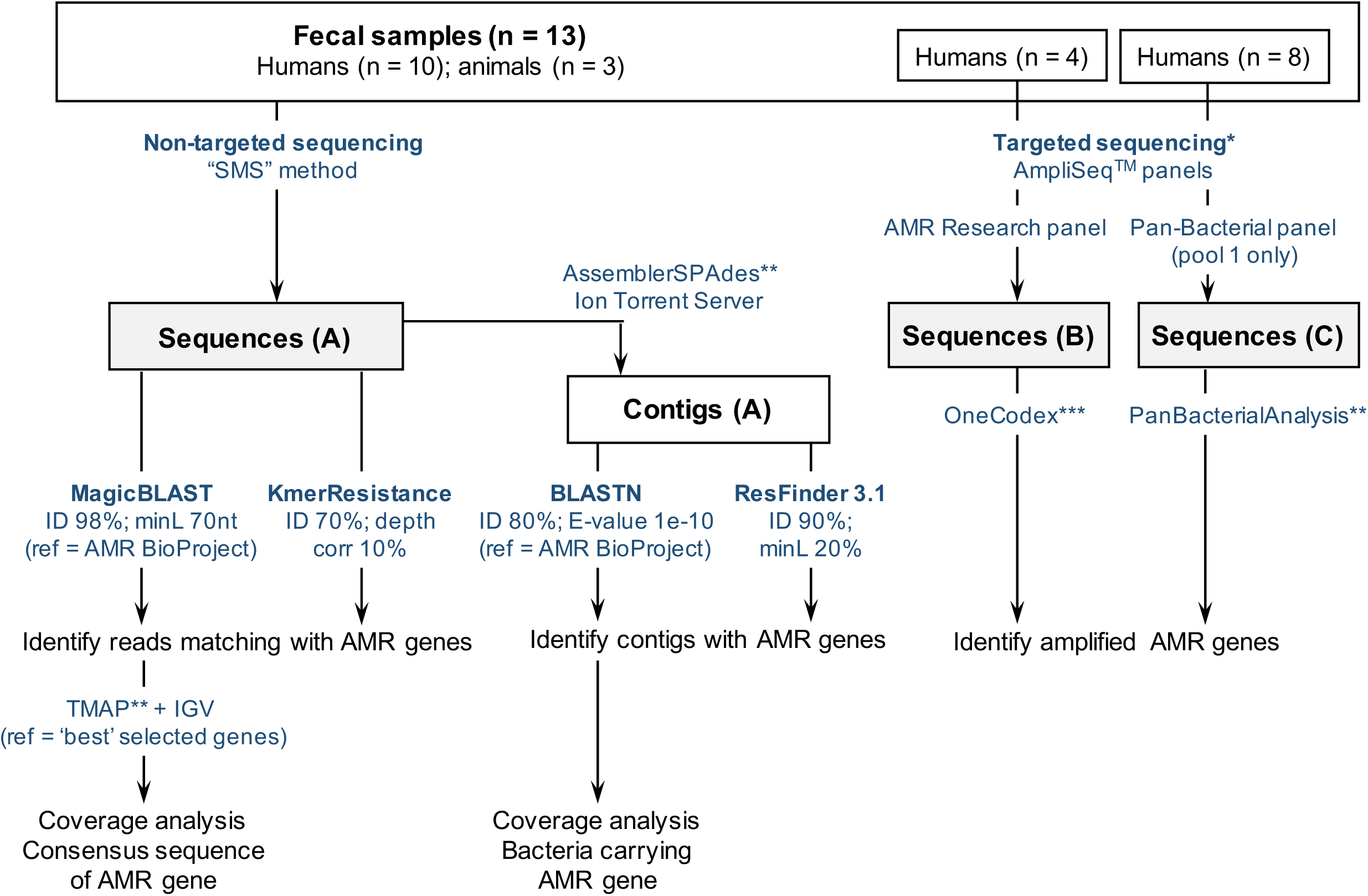
Flowchart describing three methods to identify the presence of the most common ARGs in clinically healthy human/animal samples using NGS without culturing. (A) Non-targeted sequencing of sheared DNA or “Shotgun Metagenomic Sequencing” (SMS) method, versus (B) and (C) targeted sequencing using two commercially available AmpliSeq™ panels targeting ARGs. Sequences obtained with the SMS method were analysed using different web-based tools (ResFinder and KmerResistance, https://cge.cbs.dtu.dk/services/), and an in-house method based on the comparison of reads and contigs with a NCBI-curated ARGs database (https://www.ncbi.nlm.nih.gov/bioproject/PRJNA313047). *Targeted sequencing was tested on human samples only: 4 with the AMR Research panel, 8 with the Pan-Bacterial panel. ** refers to Ion Torrent Suite plugins available from the Thermo Fisher Scientific Plugin store. *** refers to the “One Codex” online platform (available at: https://app.onecodex.com/) used with the AMR Research panel. “ID”: percentage identity threshold; “minL”: minimum length: minimum percentage of coverage compared to reference length or minimum number of nucleotides (nt); “ref”: reference sequences used for comparison with reads and/or contigs.

#### Analysis of the sequences obtained by Shotgun Metagenomic Sequencing

Raw sequences were analysed using the Ion Torrent Server linked to the Ion Torrent S5xl instrument, based on each sample having a unique barcode. We developed an “in-house method” based on the comparison of our data with references in the NCBI-curated AMR gene database (https://www.ncbi.nlm.nih.gov/bioproject/PRJNA313047) containing a total of 4,528 sequences (downloaded on the 2nd of August 2018).

The reads were queried against the NCBI’s ARGs database using Magic-BLAST^33^ with parameters as follows: 98% minimum identity between the read and the reference, and 70 nucleotides minimum length. Concomitantly, the Torrent Suite AssemblerSPAdes 5.6.0 plugin was used to generate contigs from the reads for each sample^34^. These contigs were also queried against the NCBI’s ARGs database but using nucleotide BLAST (BLASTN)^35,36^ with parameters as follows: 80% minimum identity between the contig and the reference and 1e-10 minimum E-value. The contigs identified to carry ARGs were queried against the “Microbes” reference database using BLAST Genomes in order to tentatively identify the bacterial species carrying ARGs. After identifying the ARGs present, FASTA files were created by adding chromosomes X, 21 and mitochondrion from the human genome reference sequence (hg19) to each pre-selected ARG sequence (listed in Supplementary Table S1). This was done in order to artificially increase the length of the reference used for the mapping, as it is our experience that mapping issues may arise from mapping reads against small references such as e.g. ARGs of only around 1,000 to 2,000 bp due to the scaling factor algorithm used by the Torrent Suite TMAP plugin. The extended FASTA references were added to the Ion Torrent Server and used for the mapping of sequence reads using the Torrent Suite TMAP plugin. Mapped reads and consensus sequences were visualised and analysed using the Integrative Genomics Viewer (IGV)^37^.

#### ARGs coverage and depth

Individual ARGs coverage and depth metrics were calculated using the Torrent Suite CoverageAnalysis v5.6.0.1 plugin using the filters: library type = whole genome; minimum aligned length = 100; minimum mapping quality = 20. The “average base read depth” obtained as an output was used to standardize reads counts. A “mean base depth per 5 million reads” (or BD5M) was calculated for each ARG in each sample following the formula: (average base read depth * 5,000,000) / total reads. Contrary to the common RPKM (number of reads per kb of gene per million sequenced reads), BD5M is ‘per base’ and not ‘per read’ and its interpretation is independent of the size of the reads or the reference. As an example, if 5 million reads were obtained for a sample, BD5M = 2 for a specific gene is equivalent to reads mapping the whole gene twice or reads mapping half of the gene four times. To avoid confusion between the raw coverage from the sequencer and our calculated BD5M, we hereafter refer to the latest as an abundance rather than a coverage.

#### Additional AMR analyses

We analysed the sequences obtained with the SMS method using two web-based tools, KmerResistance (available at https://cge.cbs.dtu.dk/services/KmerResistance/)^38,39^ and ResFinder 3.1 (available at https://cge.cbs.dtu.dk/services/ResFinder/)^27^, in order to compare the results with those obtained with our “in-house method” (Figure 1). Both tools rely on the same reference database (available at https://cge.cbs.dtu.dk/services/data.php)^27^. Raw sequences (FASTQ reads) from each sample were screened for ARGs present in the ResFinder database with the default settings of 70% identity threshold and 10% depth correction using “bacteria” as the host database and “resistance genes” as the gene database. Scoring method was “species determination on maximum query coverage”. Contigs (FASTA files) from each sample were also screened for ARGs present in the ResFinder database using the settings of 90% minimum identity and 20% minimum length.

#### Microbiota analysis and host identification

The Torrent Suite Ion Reporter™ software was used to process reads for metagenomics analysis of the gut microbiota. Data files (Binary Alignment Map, BAM) generated by the Ion Torrent S5xl with the SMS method were uploaded to Ion Reporter (https://ionreporter.thermofisher.com/ir/). All analyses were run using a “150-90-3” setting, i.e. minimum read length = 150; minimum alignment coverage between hit and query = 90%; minimum number of unique reads (or ‘read abundance filter’) = 3. This means that only reads found as triplicates or more were included in the analysis. A minimum percentage identity of 97% was required for genus identification of a ribosomal sequence, and a minimum percentage identity of 99% for species identification. When there was a difference of less than 0.2% match between the top hit and the next best hit, the sequence was identified as ‘species slash ID’ (unidentified species). The relative abundance of the species/genus identified in the gut microbiota was determined from Ion Reporter™.

The contigs generated from SMS allowed tentative identification of host bacteria carrying the ARGs. This was achieved by running the contigs against the “Microbes” database using BLAST Genomes or the ResFinder tool.

#### Analysis of the sequences obtained by Ion AmpliSeq™ panels

Following targeted amplification, sequences obtained from the Ion Torrent S5xl were analysed based on each sample having a unique barcode and using dedicated platforms or plugins, i.e. (i) the “One Codex” web-based platform (available at: https://app.onecodex.com/) for the sequences obtained from using the AMR Research panel, or (ii) the “PanBacterialAnalysis” Torrent Suite plugin from the Ion Torrent Server for the sequences obtained from using the Pan-Bacterial Research panel (see Figure 1). One Codex reported the percentage of the ARG sequence covered by reads in the sample, the percentage identity to the reference ARG sequence, and the depth of read coverage. The “PanBacterialAnalysis” plugin reported the “read counts” matching each ARG in the database.

## Results

All 13 samples were processed using the SMS method; with eight samples processed using the Pan-Bacterial Research panel, and four of these eight also processed using the AMR panel (Table 2). SMS generated a total of 3.8 to 11 million reads per sample, with average read lengths of 162-201 nucleotides (Table 2). Whatever the method, all samples had reads that mapped to ARGs, with abundance and composition varying between samples. SMS overall identified ARGs to 2-11 classes of antibiotics in individual samples (Figure 2 and Table 3), with ARGs to two additional classes identified using targeted sequencing methods.

**Table 2.** Summary of NGS sequencing results using three methods: (1) shotgun metagenomics sequencing (SMS) or (2) targeted sequencing using community panels from Life Technologies: (2a) the Ion AmpliSeq™ Antimicrobial Resistance (AMR) Research panel and (2b) the Ion AmpliSeq™ Pan-Bacterial Research (PBR) panel. BC: barcode adapter identifier; Reads: number of short read sequences per sample; MRL: mean read length in nucleotides. The four human samples tested with all three methods are in dark grey, the four human samples tested with two methods are in light grey and the five samples tested only with the SMS method are in white.

| Sample ID | Host | SMS method |  |  | AMR panel |  | PBR panel |  |
| --- | --- | --- | --- | --- | --- | --- | --- | --- |
|  |  | BC | Reads | MRL | BC | Reads | BC | Reads |
| ST4-3mo | Human - child | BC062 | 5,057,157 | 184 | - | - | - | - |
| ST5-1mo | Human - child | BC002 | 10,998,563 | 201 | BC049 | 1,486,710 | BC053 | 505,192 |
| ST5-18mo | Human - child | BC063 | 5,656,858 | 195 | - | - | - | - |
| HS21 | Human - adult | BC021 | 3,829,581 | 182 | BC050 | 523,118 | BC054 | 524,591 |
| HS22 | Human - adult | BC022 | 5,721,672 | 184 | - | - | BC055 | 377,819 |
| HS23 | Human - adult | BC023 | 9,313,335 | 197 | - | - | BC056 | 262,060 |
| HS24 | Human - adult | BC024 | 7,303,016 | 195 | BC051 | 709,047 | BC057 | 491,234 |
| HS25 | Human - adult | BC025 | 6,986,354 | 193 | - | - | BC058 | 174,895 |
| HS26 | Human - adult | BC026 | 9,411,554 | 198 | BC052 | 730,272 | BC059 | 431,487 |
| HS28 | Human - adult | BC028 | 7,952,415 | 180 | - | - | BC060 | 110,975 |
| ST1-MAD | Pacific black ducks | BC065 | 5,671,145 | 162 | - | - | - | - |
| ST3-MUD | Muscovy duck | BC066 | 8,438,563 | 169 | - | - | - | - |
| DFS | Pool from 2 dogs | BC070 | 8,570,346 | 192 | - | - | - | - |

**Table 3.**
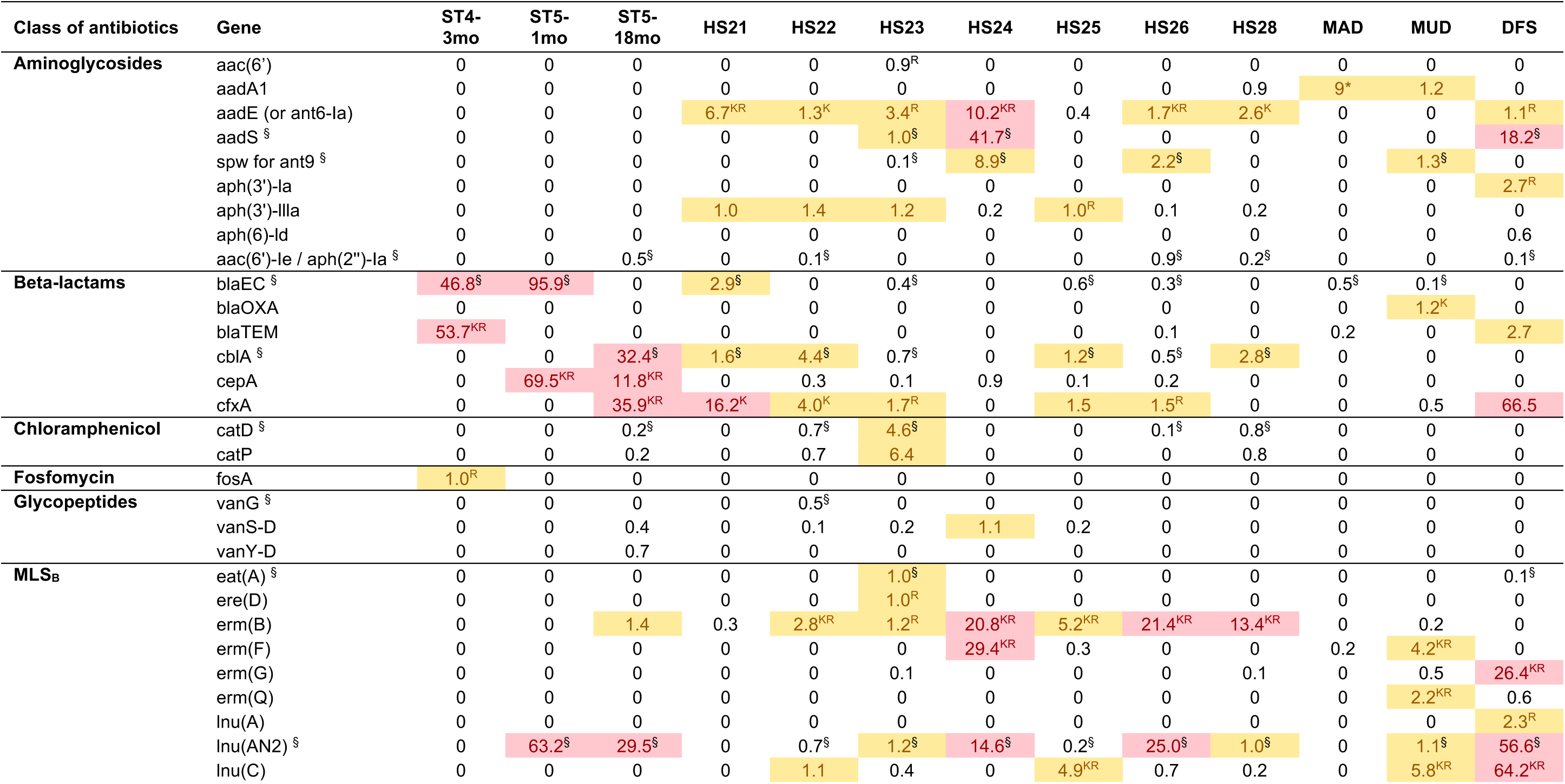
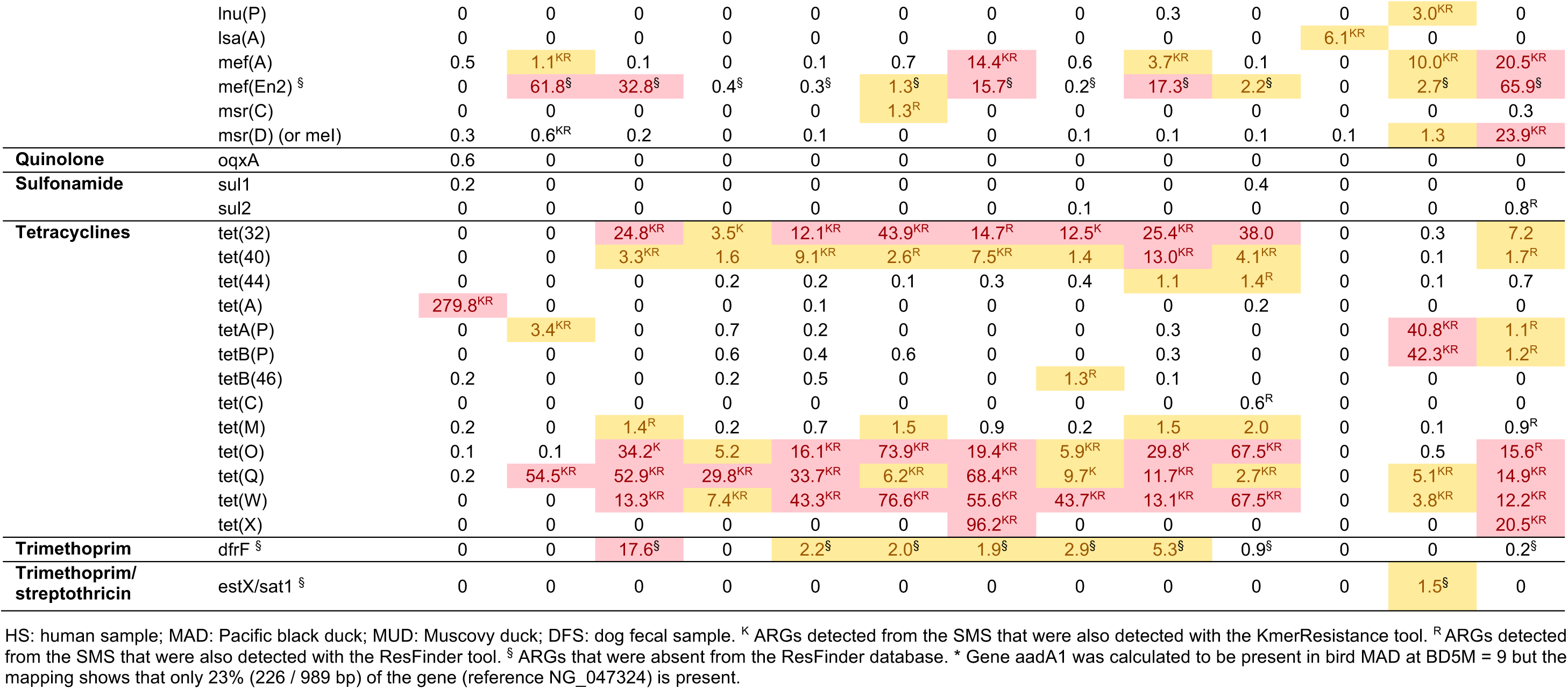
Antimicrobial resistance genes (ARGs) identified from the shotgun metagenomics sequencing (SMS) using the “in-house” analysis. (based on queries against the NCBI’s “Bacterial Antimicrobial Resistance Reference Gene Database” BioProject PRJNA313047). Mean depth coverage per 5 million reads (BD5M) between 1 and 10 is highlighted in yellow, BD5M > 10 is highlighted in red. Additional ARGs identified with the ResFinder and/or KmerResistance tools but not the in-house method are not listed here (see text).

| Class of antibiotics | Gene | ST4-3mo | ST5-1mo | ST5-18mo | HS21 | HS22 | HS23 | HS24 | HS25 | HS26 | HS28 | MAD | MUD | DFS |
| --- | --- | --- | --- | --- | --- | --- | --- | --- | --- | --- | --- | --- | --- | --- |
| **Aminoglycosides** | aac(6') | 0 | 0 | 0 | 0 | 0 | 0.9<sup>R</sup> | 0 | 0 | 0 | 0 | 0 | 0 | 0 |
|  | aadA1 | 0 | 0 | 0 | 0 | 0 | 0 | 0 | 0 | 0 | 0.9 | 9* | 1.2 | 0 |
|  | aadE (or ant6-la) | 0 | 0 | 0 | 6.7<sup>KR</sup> | 1.3<sup>K</sup> | 3.4<sup>R</sup> | 10.2<sup>KR</sup> | 0.4 | 1.7<sup>KR</sup> | 2.6<sup>K</sup> | 0 | 0 | 1.1<sup>R</sup> |
|  | aadS <sup>§</sup> | 0 | 0 | 0 | 0 | 0 | 1.0<sup>§</sup> | 41.7<sup>§</sup> | 0 | 0 | 0 | 0 | 0 | 18.2<sup>§</sup> |
|  | spw for ant9 <sup>§</sup> | 0 | 0 | 0 | 0 | 0 | 0.1<sup>§</sup> | 8.9<sup>§</sup> | 0 | 2.2<sup>§</sup> | 0 | 0 | 1.3<sup>§</sup> | 0 |
|  | aph(3')-la | 0 | 0 | 0 | 0 | 0 | 0 | 0 | 0 | 0 | 0 | 0 | 0 | 2.7<sup>R</sup> |
|  | aph(3')-IIIa | 0 | 0 | 0 | 1.0 | 1.4 | 1.2 | 0.2 | 1.0<sup>R</sup> | 0.1 | 0.2 | 0 | 0 | 0 |
|  | aph(6)-Id | 0 | 0 | 0 | 0 | 0 | 0 | 0 | 0 | 0 | 0 | 0 | 0 | 0.6 |
|  | aac(6')-Ie / aph(2'')-Ia <sup>§</sup> | 0 | 0 | 0.5<sup>§</sup> | 0 | 0.1<sup>§</sup> | 0 | 0 | 0 | 0.9<sup>§</sup> | 0.2<sup>§</sup> | 0 | 0 | 0.1<sup>§</sup> |
| **Beta-lactams** | blaEC <sup>§</sup> | 46.8<sup>§</sup> | 95.9<sup>§</sup> | 0 | 2.9<sup>§</sup> | 0 | 0.4<sup>§</sup> | 0 | 0.6<sup>§</sup> | 0.3<sup>§</sup> | 0 | 0.5<sup>§</sup> | 0.1<sup>§</sup> | 0 |
|  | blaOXA | 0 | 0 | 0 | 0 | 0 | 0 | 0 | 0 | 0 | 0 | 0 | 1.2<sup>K</sup> | 0 |
|  | blaTEM | 53.7<sup>KR</sup> | 0 | 0 | 0 | 0 | 0 | 0 | 0 | 0.1 | 0 | 0.2 | 0 | 2.7 |
|  | cbIA <sup>§</sup> | 0 | 0 | 32.4<sup>§</sup> | 1.6<sup>§</sup> | 4.4<sup>§</sup> | 0.7<sup>§</sup> | 0 | 1.2<sup>§</sup> | 0.5<sup>§</sup> | 2.8<sup>§</sup> | 0 | 0 | 0 |
|  | cepA | 0 | 69.5<sup>KR</sup> | 11.8<sup>KR</sup> | 0 | 0.3 | 0.1 | 0.9 | 0.1 | 0.2 | 0 | 0 | 0 | 0 |
|  | cfxA | 0 | 0 | 35.9<sup>KR</sup> | 16.2<sup>K</sup> | 4.0<sup>K</sup> | 1.7<sup>R</sup> | 0 | 1.5 | 1.5<sup>R</sup> | 0 | 0 | 0.5 | 66.5 |
| **Chloramphenicol** | catD <sup>§</sup> | 0 | 0 | 0.2<sup>§</sup> | 0 | 0.7<sup>§</sup> | 4.6<sup>§</sup> | 0 | 0 | 0.1<sup>§</sup> | 0.8<sup>§</sup> | 0 | 0 | 0 |
|  | catP | 0 | 0 | 0.2 | 0 | 0.7 | 6.4 | 0 | 0 | 0 | 0.8 | 0 | 0 | 0 |
| **Fosfomycin** | fosA | 1.0<sup>R</sup> | 0 | 0 | 0 | 0 | 0 | 0 | 0 | 0 | 0 | 0 | 0 | 0 |
| **Glycopeptides** | vanG <sup>§</sup> | 0 | 0 | 0 | 0 | 0.5<sup>§</sup> | 0 | 0 | 0 | 0 | 0 | 0 | 0 | 0 |
|  | vanS-D | 0 | 0 | 0.4 | 0 | 0.1 | 0.2 | 1.1 | 0.2 | 0 | 0 | 0 | 0 | 0 |
|  | vanY-D | 0 | 0 | 0.7 | 0 | 0 | 0 | 0 | 0 | 0 | 0 | 0 | 0 | 0 |
| **MLS<sub>B</sub>** | eat(A) <sup>§</sup> | 0 | 0 | 0 | 0 | 0 | 1.0<sup>§</sup> | 0 | 0 | 0 | 0 | 0 | 0 | 0.1<sup>§</sup> |
|  | ere(D) | 0 | 0 | 0 | 0 | 0 | 1.0<sup>R</sup> | 0 | 0 | 0 | 0 | 0 | 0 | 0 |
|  | erm(B) | 0 | 0 | 1.4 | 0.3 | 2.8<sup>KR</sup> | 1.2<sup>R</sup> | 20.8<sup>KR</sup> | 5.2<sup>KR</sup> | 21.4<sup>KR</sup> | 13.4<sup>KR</sup> | 0 | 0.2 | 0 |
|  | erm(F) | 0 | 0 | 0 | 0 | 0 | 0 | 29.4<sup>KR</sup> | 0.3 | 0 | 0 | 0.2 | 4.2<sup>KR</sup> | 0 |
|  | erm(G) | 0 | 0 | 0 | 0 | 0 | 0.1 | 0 | 0 | 0 | 0.1 | 0 | 0.5 | 26.4<sup>KR</sup> |
|  | erm(Q) | 0 | 0 | 0 | 0 | 0 | 0 | 0 | 0 | 0 | 0 | 0 | 2.2<sup>KR</sup> | 0.6 |
|  | lnu(A) | 0 | 0 | 0 | 0 | 0 | 0 | 0 | 0 | 0 | 0 | 0 | 0 | 2.3<sup>R</sup> |
|  | lnu(AN2) <sup>§</sup> | 0 | 63.2<sup>§</sup> | 29.5<sup>§</sup> | 0 | 0.7<sup>§</sup> | 1.2<sup>§</sup> | 14.6<sup>§</sup> | 0.2<sup>§</sup> | 25.0<sup>§</sup> | 1.0<sup>§</sup> | 0 | 1.1<sup>§</sup> | 56.6<sup>§</sup> |
|  | lnu(C) | 0 | 0 | 0 | 0 | 1.1 | 0.4 | 0 | 4.9<sup>KR</sup> | 0.7 | 0.2 | 0 | 5.8<sup>KR</sup> | 64.2<sup>KR</sup> |
|  | lnu(P) | 0 | 0 | 0 | 0 | 0 | 0 | 0 | 0 | 0.3 | 0 | 0 | 3.0<sup>KR</sup> | 0 |
|  | lsa(A) | 0 | 0 | 0 | 0 | 0 | 0 | 0 | 0 | 0 | 0 | 6.1<sup>KR</sup> | 0 | 0 |
|  | mef(A) | 0.5 | 1.1<sup>KR</sup> | 0.1 | 0 | 0.1 | 0.7 | 14.4<sup>KR</sup> | 0.6 | 3.7<sup>KR</sup> | 0.1 | 0 | 10.0<sup>KR</sup> | 20.5<sup>KR</sup> |
|  | mef(En2) <sup>§</sup> | 0 | 61.8<sup>§</sup> | 32.8<sup>§</sup> | 0.4<sup>§</sup> | 0.3<sup>§</sup> | 1.3<sup>§</sup> | 15.7<sup>§</sup> | 0.2<sup>§</sup> | 17.3<sup>§</sup> | 2.2<sup>§</sup> | 0 | 2.7<sup>§</sup> | 65.9<sup>§</sup> |
|  | msr(C) | 0 | 0 | 0 | 0 | 0 | 1.3<sup>R</sup> | 0 | 0 | 0 | 0 | 0 | 0 | 0.3 |
|  | msr(D) (or mel) | 0.3 | 0.6<sup>KR</sup> | 0.2 | 0 | 0.1 | 0 | 0 | 0.1 | 0.1 | 0.1 | 0.1 | 1.3 | 23.9<sup>KR</sup> |
| **Quinolone** | oqxA | 0.6 | 0 | 0 | 0 | 0 | 0 | 0 | 0 | 0 | 0 | 0 | 0 | 0 |
| **Sulfonamide** | sul1 | 0.2 | 0 | 0 | 0 | 0 | 0 | 0 | 0 | 0 | 0.4 | 0 | 0 | 0 |
|  | sul2 | 0 | 0 | 0 | 0 | 0 | 0 | 0 | 0.1 | 0 | 0 | 0 | 0 | 0.8<sup>R</sup> |
| **Tetracyclines** | tet(32) | 0 | 0 | 24.8<sup>KR</sup> | 3.5<sup>K</sup> | 12.1<sup>KR</sup> | 43.9<sup>KR</sup> | 14.7<sup>R</sup> | 12.5<sup>K</sup> | 25.4<sup>KR</sup> | 38.0 | 0 | 0.3 | 7.2 |
|  | tet(40) | 0 | 0 | 3.3<sup>KR</sup> | 1.6 | 9.1<sup>KR</sup> | 2.6<sup>R</sup> | 7.5<sup>KR</sup> | 1.4 | 13.0<sup>KR</sup> | 4.1<sup>KR</sup> | 0 | 0.1 | 1.7<sup>R</sup> |
|  | tet(44) | 0 | 0 | 0 | 0.2 | 0.2 | 0.1 | 0.3 | 0.4 | 1.1 | 1.4<sup>R</sup> | 0 | 0.1 | 0.7 |
|  | tet(A) | 279.8<sup>KR</sup> | 0 | 0 | 0 | 0.1 | 0 | 0 | 0 | 0 | 0.2 | 0 | 0 | 0 |
|  | tetA(P) | 0 | 3.4<sup>KR</sup> | 0 | 0.7 | 0.2 | 0 | 0 | 0 | 0.3 | 0 | 0 | 40.8<sup>KR</sup> | 1.1<sup>R</sup> |
|  | tetB(P) | 0 | 0 | 0 | 0.6 | 0.4 | 0.6 | 0 | 0 | 0.3 | 0 | 0 | 42.3<sup>KR</sup> | 1.2<sup>R</sup> |
|  | tetB(46) | 0.2 | 0 | 0 | 0.2 | 0.5 | 0 | 0 | 1.3<sup>R</sup> | 0.1 | 0 | 0 | 0 | 0 |
|  | tet(C) | 0 | 0 | 0 | 0 | 0 | 0 | 0 | 0 | 0 | 0.6<sup>R</sup> | 0 | 0 | 0 |
|  | tet(M) | 0.2 | 0 | 1.4<sup>R</sup> | 0.2 | 0.7 | 1.5 | 0.9 | 0.2 | 1.5 | 2.0 | 0 | 0.1 | 0.9<sup>R</sup> |
|  | tet(O) | 0.1 | 0.1 | 34.2<sup>K</sup> | 5.2 | 16.1<sup>KR</sup> | 73.9<sup>KR</sup> | 19.4<sup>KR</sup> | 5.9<sup>KR</sup> | 29.8<sup>K</sup> | 67.5<sup>KR</sup> | 0 | 0.5 | 15.6<sup>R</sup> |
|  | tet(Q) | 0.2 | 54.5<sup>KR</sup> | 52.9<sup>KR</sup> | 29.8<sup>KR</sup> | 33.7<sup>KR</sup> | 6.2<sup>KR</sup> | 68.4<sup>KR</sup> | 9.7<sup>K</sup> | 11.7<sup>KR</sup> | 2.7<sup>KR</sup> | 0 | 5.1<sup>KR</sup> | 14.9<sup>KR</sup> |
|  | tet(W) | 0 | 0 | 13.3<sup>KR</sup> | 7.4<sup>KR</sup> | 43.3<sup>KR</sup> | 76.6<sup>KR</sup> | 55.6<sup>KR</sup> | 43.7<sup>KR</sup> | 13.1<sup>KR</sup> | 67.5<sup>KR</sup> | 0 | 3.8<sup>KR</sup> | 12.2<sup>KR</sup> |
|  | tet(X) | 0 | 0 | 0 | 0 | 0 | 0 | 96.2<sup>KR</sup> | 0 | 0 | 0 | 0 | 0 | 20.5<sup>KR</sup> |
| **Trimethoprim** | dfrF <sup>§</sup> | 0 | 0 | 17.6<sup>§</sup> | 0 | 2.2<sup>§</sup> | 2.0<sup>§</sup> | 1.9<sup>§</sup> | 2.9<sup>§</sup> | 5.3<sup>§</sup> | 0.9<sup>§</sup> | 0 | 0 | 0.2<sup>§</sup> |
| **Trimethoprim/**<br>**streptothricin** | estX/sat1 <sup>§</sup> | 0 | 0 | 0 | 0 | 0 | 0 | 0 | 0 | 0 | 0 | 0 | 1.5<sup>§</sup> | 0 |

HS: human sample; MAD: Pacific black duck; MUD: Muscovy duck; DFS: dog fecal sample. <sup>K</sup> ARGs detected from the SMS that were also detected with the KmerResistance tool. <sup>R</sup> ARGs detected from the SMS that were also detected with the ResFinder tool. <sup>§</sup> ARGs that were absent from the ResFinder database. \* Gene aadA1 was calculated to be present in bird MAD at BD5M = 9 but the mapping shows that only 23% (226 / 989 bp) of the gene (reference NG\_047324) is present.

**Figure 2.**
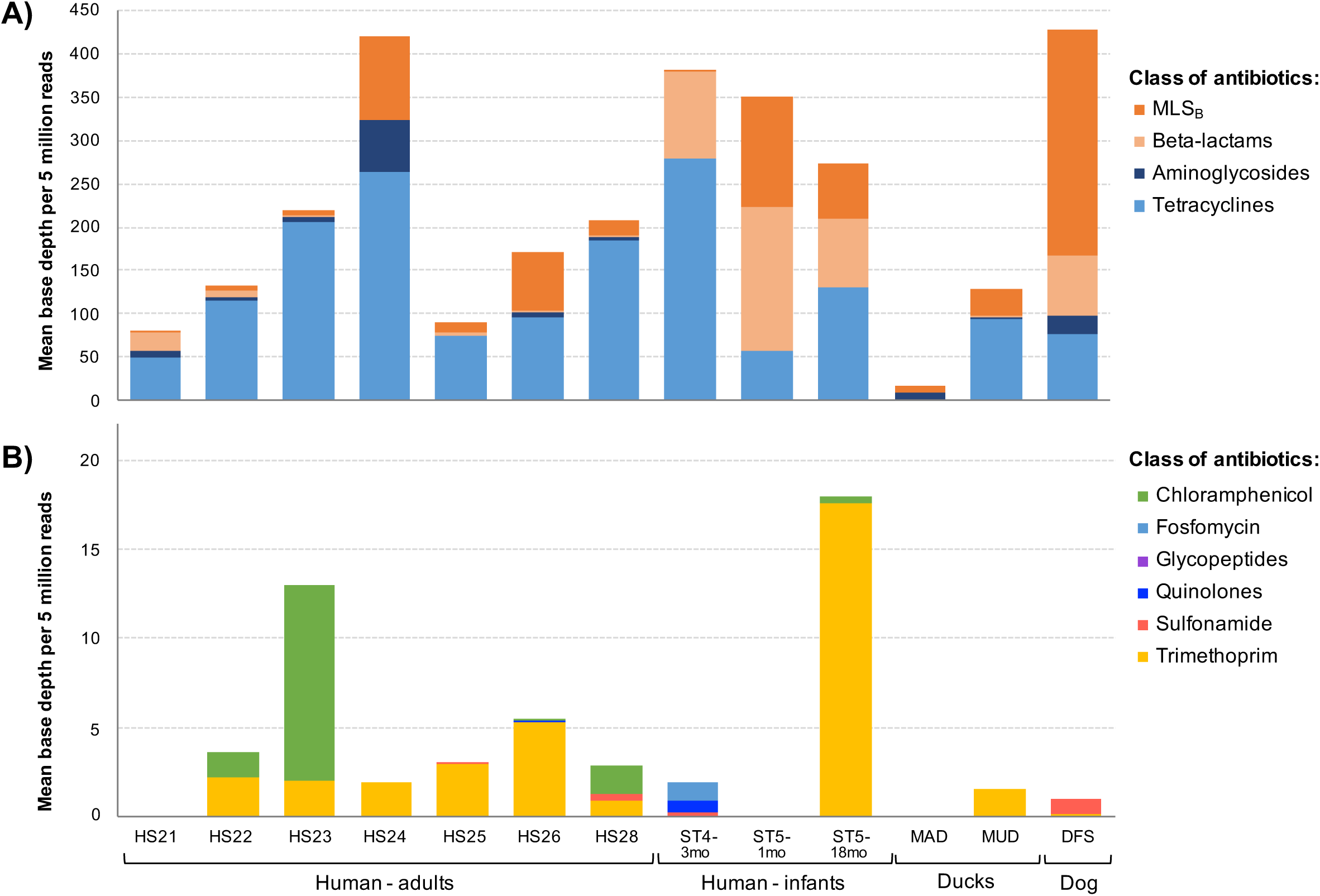
Overview of antimicrobial resistance genes (ARGs) abundance and composition in human and animal samples (n = 13) using the shotgun metagenomics sequencing (SMS) method. (A) Major *versus* (B) minor ARGs. ARGs abundance is calculated as a “mean base depth per 5 million reads” (BD5M) for each reference gene in each sample, and then summed for ARGs belonging to the same class of antibiotics.

### SMS-based analysis of ARGs

All 13 samples were processed using the SMS method, allowing comparison of ARGs diversity between the human samples; three young infants and seven adults; as well as the pooled domestic dog fecal sample (DFS), and two bird samples (MAD being a pool of six birds). We identified ARGs from both the reads and the contigs (see Table S1).

#### Resistance - antibiotic classes

Using the “in-house method”, we detected genes conferring resistance to 11 classes of antibiotics (out of 16 classes included in the NCBI’s ARGs database). ARGs conferring resistance to at least one class of antibiotics at BD5M >50 were found in 12 samples (Figure 2 and Table 3). ARGs conferring resistance to tetracycline (BD5M 49 to 280), macrolide, lincosamide, streptogramin B (MLS_B_) (BD5M 0.7 to 260), β-lactams (BD5M 1.9 to 165) and aminoglycoside (BD5M up to 61) were found at higher abundances in all 13 samples (Figure 2A, Table 3). ARGs to six other classes of antibiotics were found in ten samples at lower abundance (Figure 2B): trimethoprim (BD5M up to 17.6), chloramphenicol (BD5M up to 11), glycopeptides, quinolones, fosfomycin and sulfonamide (all last four classes with BD5M ≤ 1) (Figure 2B and Table 3). An eleventh class of antibiotics was only detected as a bifunctional gene estX/sat1 conferring resistance to trimethoprim/streptothricin.

The wild ducks (MAD) and the 1-month old human (ST5-1mo) carried genes conferring resistance to the fewest classes of antibiotics –three classes (or two classes at BD5M ≥ 1 for sample MAD)– while HS23, HS24, duck MUD and dog DFS carried ARGs attributable to the most classes of antibiotics, i.e. seven classes (or six classes at BD5M ≥ 1) (Table 3).

#### Resistance - gene level (ARGs)

At the gene level, 62 ARGs were detected from the SMS-generated reads: 54 using the “in-house method” (or 45 at BD5M ≥ 1), 43 using the ResFinder tool and 28 using the KmerResistance tool (Figure 3B). Within a same class of antibiotics, the lowest diversity of ARGs was observed for fosfomycin, quinolones and trimethoprim/streptothricin (n = 1), while the highest diversity of ARGs was observed for MLS_B_ (n = 15) and tetracycline (n = 13) (Table 3).

**Figure 3.**
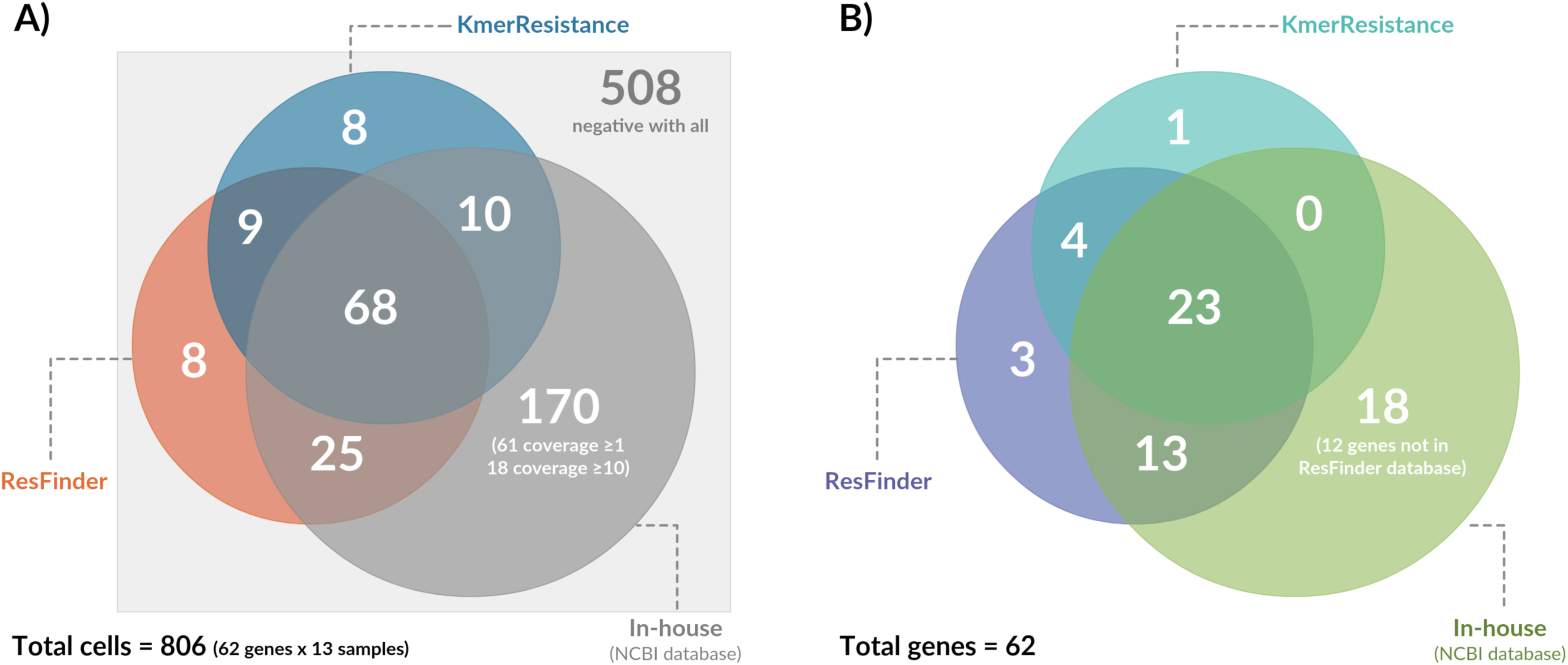
Resistance patterns identified from shotgun metagenomics sequencing (SMS) using three analysis methods (A) per gene and per sample or (B) per gene. SMS reads were screened for the presence of antimicrobial resistance genes using (i) an “in-house method” (based on queries against the NCBI’s AMR reference gene database BioProject PRJNA313047), (ii) ResFinder and/or (iii) KmerResistance tools.

Out of 54 ARGs identified with the in-house method, 18 could not be detected using the web based tools. Of these, twelve were absent from the ResFinder database and thus could not have been detected using the web-based tools (Figure 3B). Conversely, eight ARGs identified from the SMS reads using the ResFinder and KmerResistance tools could not be detected using the in-house method (Figure 3B).

#### Incongruent results between methods at the individual level

ARGs identified by all three methods at the gene level could sometimes be identified by only two or one method at the individual level (see supplementary Table S2 for detailed results).

Gene cfxA (β-lactam resistance) was detected with all three methods in sample ST5-18mo but was detected only with the in-house method in dog DFS (BD5M = 66.5). The best match in the NCBI’s AMR reference database (NG_047636.1) was only 89% similar to the consensus sequence, while the best match by BLASTN against the full NCBI nucleotide database (98.9% similarity) was gene cfxA6 (GQ342996), which was absent from the NCBI’s AMR database.

Gene tet(32) (tetracycline resistance) was detected only with the in-house method in HS28 (BD5M = 38.0) with the best match in the NCBI’s AMR reference database being NG_048124.1. However, reads coverage against NG_048124.1 was very high for the first 350 bp and last 700 bp only, probably related to recombination events with other ‘tet’ genes (see Discussion). The best match with BLASTN (97.8% similarity) was a recombinant tet(O/32/O) gene (JQ740052). Even though the ResFinder database includes five recombinant tet(O/32/O) gene references, it was not detected in HS28 with the web-based tools.

At the individual level, ARGs identified using the web-based tools but not the in-house method were: aadA24 and aph(3’)-III (aminoglycosides); cfxA6 (β-lactams); catS (chloramphenicol); mdf(A) (MLS_B_); tet(O/W), tet(O/32/O) and tet(W/32/O) (tetracycline). Out of these, seven genes were absent from the NCBI’s AMR database, and thus could not have been detected using the in-house method (Table S2). The remaining gene (aadA24) was detected only with the web-based tools despite being present in both reference databases, but further analysis showed that this gene was detected as an aadA1 gene with the in-house method.

#### Resistance profiles between and within species

Different species (human / birds / dogs) and different samples showed different resistance profiles (Table 3). With the in-house method, HS23 showed the highest AMR diversity within the human samples, with 30 ARGs (or 19 at BD5M ≥ 1) belonging to six classes of antibiotics detected, followed by HS26 (28 ARGs or 15 at BD5M ≥ 1), and HS22 (28 ARGs or 12 at BD5M ≥ 1). HS24 showed the highest abundances of individual ARGs, with 12 out of 20 ARGs detected at BD5M ≥ 10. When accounting for the additional genes detected using the ResFinder and KmerResistance tools, the number of ARGs increased for most samples, HS22 becoming the sample with the highest AMR diversity (34 ARGs) (Table S2).

Most of the adult samples were similar to each other in their resistance profile, except for HS24 (Figure 2). The three infants showed higher abundances for fewer ARGs compared to the adults, e.g. three ARGs at BD5M ≥ 1 in infant ST4-3mo: two ARGs conferring resistance to β-lactams (BD5M = 47 and 54) and one to tetracycline (BD5M = 280) (Table 3).

The sample MAD (pool from 6 Pacific black ducks) had the lowest AMR diversity, with two ARGs at BD5M ≥ 1 (BD5M = 9 for aminoglycosides; 6.4 for MLS_B_), while the sample MUD (from a Muscovy duck) showed the presence of 16 ARGs at BD5M ≥ 1 (resistance to aminoglycoside, MLS_B_, β-lactams and tetracycline). ARGs in highest abundance in the sample MUD were tetA(P) and tetB(P) (tetracycline resistance; BD5M > 40), two genes also found in humans and dog DFS but at very low levels (Table 3).

The pooled DFS showed one of the highest AMR diversity, with 29 ARGs detected (or 20 at BD5M ≥ 1) belonging to six classes of antibiotics; this number increased to 32 ARGs when accounting for additional genes detected using the ResFinder and KmerResistance tools. The dog sample also contained 12 ARGs at high abundance (BD5M ≥ 10), including 6 MLS_B_ genes. This is comparable to 5 MLS_B_ genes at BD5M ≥ 10 found in HS24, but with only three ARGs in common, and those in the dog DFS typically at higher abundances than in the human HS24.

### Identification of bacteria carrying ARGs

The metagenomics analysis of the gut microbiome for selected samples (n = 7) using the Torrent Suite Ion Reporter™ software is summarized in Table 4. Because adult human microbiotas are diverse and complex, we are detailing one example for which the ARGs carrying bacteria were likely identified (HS24).

**Table 4.** Percentages of major bacterial Orders in the fecal microbiome of selected samples (n = 7). Bacterial orders likely carrying antimicrobial resistance genes (ARGs) identified in each sample are highlighted as a grey box.

| Phylum | Order | ST5-1mo | ST5-18mo | ST4-3mo | HS24 | DFS | ST1-MAD | ST3-MUD |
| --- | --- | --- | --- | --- | --- | --- | --- | --- |
| Firmicutes | Clostridiales | < 1% | 35% |  | 41% | 4% | 9% | 54% |
|  | Erysipelotrichales |  |  |  |  |  | 4% | 18% |
|  | Lactobacillales | < 1% |  |  |  | 42% | 74% |  |
|  | Selenomonadales |  |  |  |  | 12% |  |  |
|  | Veillonellales |  |  | 2% |  |  |  | 2% |
| Bacteroidetes | Bacteroidales | 34% | 51% |  | 56% | 30% |  |  |
| Proteobacteria | Burkholderiales |  | 5% |  |  |  |  |  |
|  | Campylobacteriales |  |  |  |  |  | 2% | 19% |
|  | Enterobacteriales | 66% |  | 46% |  |  |  |  |
|  | Gammaproteobacteria |  |  |  |  | 5% |  |  |
|  | Kopriimonadales |  |  |  |  |  | 1% |  |
|  | Legionellales |  |  |  |  |  |  | 2% |
| Verrucomicrobia | Verrucomicrobiales |  | 6% |  | 2% |  |  |  |
| Actinobacteria | Bifidobacteriales |  | 1% | 51% |  |  |  |  |
| Cyanobacteria | Synechococcales |  |  |  |  |  | 5% |  |
|  | Nostocales |  |  |  |  |  | 3% |  |
| Deinococcus-Thermus | Deinococcales |  |  |  |  |  | 1% |  |

ST5-1mo: infant sample one month old; ST5-18mo: infant sample 18 months old; ST4-3mo: infant sample three months old; HS: human sample; DFS: dog fecal sample; MAD: pool from 6 Pacific black ducks; MUD: Muscovy duck.

#### Human infants samples

The composition of the humans gut microbiota was age-related (Table 4). At the taxonomic level ‘Order’, there were two overwhelmingly dominant bacterial populations in ST5-1mo: Enterobacteriales (66%) and Bacteroidales (34%). ARGs were identified in both populations: Order Enterobacteriales (possibly Species *Escherichia coli*) carrying the blaEC gene for cephalosporin-hydrolyzing class C β-lactamase; and Order Bacteroidales (possibly Species *Bacteroides fragilis*) carrying three ARGs, one to β-lactams, and two to MLS_B_ antibiotics.

In ST5-18mo (same child as ST5-1mo but older) Order Enterobacteriales was absent, but new Orders appeared: Bacteroidales (51%), Clostridiales (35%), Verrucomicrobiales (6%) and Burkholderiales (5%). ARGs to three β-lactams and two MLS_B_ antibiotics were most likely carried by bacteria in the Order Bacteroidales, while ARGs to tetracyclines were most likely carried by bacteria in various taxa, predominantly Order Clostridiales and Order Bacteroidales. In ST4-3mo, we detected no or little bacteria in Order Bacteroidales, but many bacteria in Bifidobacteriales (51%) and Enterobacteriales (46%). ARGs were identified in bacteria belonging to Order Enterobacteriales only: two ARGs to β-lactams, one to fosfomycin, one to MLS_B_ antibiotics and one to tetracycline. Of note, genes blaTEM and tet(A) were likely carried by a plasmid, as identified by a BLASTn query of the contigs carrying these genes.

#### Human adult samples

In sample HS24, the two main Orders were Bacteroidales (56%) and Clostridiales (41%). The most abundant ARGs appeared to be carried by bacteria in the Order Bacteroidales (resistance to aminoglycosides, MLS_B_ and tetracyclines). Of note, gene erm(B) (conferring resistance to macrolides) was likely carried by a plasmid.

#### Animal samples

The gut microbiota and resistant bacterial species in the dog sample DFS were somewhat similar to the human samples, with bacteria in the Order Bacteroidales likely carrying one β-lactam-resistance gene, while 6 MLS_B_-resistance genes were likely carried by bacteria in the Phylum Firmicutes (Order unidentified). Bacteria carrying ARGs to tetracycline and aminoglycosides could not be definitively identified. The aph(3’)-Ia gene (conferring resistance to aminoglycosides) was likely sitting on a plasmid also carrying a transposase, and the aadS gene (aminoglycosides) was likely sitting on a transposon.

The birds’ microbiotas were different from the mammals (Table 3). Bacteria from the Order Lactobacillales (possibly Species *Enterococcus faecalis*) were likely carrying gene lsa(A) (conferring multidrug resistance) detected in the wild pacific black ducks (MAD). Bacteria from the Order Clostridia (possibly Species *Clostridium perfringens*) was likely carrying genes erm(Q) (resistance to MLS_B_) and tetA(P) and tetB(P) (resistance to tetracycline) detected in the Muscovy duck (MUD), with other less abundant MLS_B_-resistance genes likely carried by unidentified bacteria (Phylum Firmicutes).

### Comparison of the SMS method and the targeted sequencing methods to detect ARGs

Four human samples were analysed using all three methods. For the SMS method, we focused on the in-house analysis and ARGs at BD5M ≥ 1. The results are summarized in Table 5 (see Table S3 for detailed information per sample and per gene).

**Table 5.** Number of antimicrobial resistance genes (ARGs) identified with three sequencing methods from human samples (n = 4). Reads obtained from the shotgun metagenomics sequencing (SMS) were queried against the NCBI’s AMR reference gene database BioProject PRJNA313047. Other methods were targeted sequencing using the Ion AmpliSeq™ AMR Research panel or the Ion AmpliSeq™ Pan-Bacterial panel.

|  | Method | ST5-1mo | HS21 | HS24 | HS26 |
| --- | --- | --- | --- | --- | --- |
| Number of ARGs identified with each method | AMR Research panel * | 29 (14) | 39 (33) | 34 (28) | 54 (44) |
|  | Pan-Bacterial panel | 14 | 26 | 23 | 30 |
|  | SMS + in-house ** | 9 (7) | 17 (10) | 20 (16) | 28 (15) |
| Number of ARGs identified with any method |  | 34 | 45 | 39 | 64 |
| Number of ARGs identified exclusively with one method | AMR Research panel only | 24 (66.7%) | 27 (55.1%) | 21 (50.0%) | 38 (55.9%) |
|  | Pan-Bacterial panel only | 2 (5.6%) | 3 (6.1%) | 3 (7.1%) | 3 (4.4%) |
|  | SMS + in-house only | 5 (13.9%) | 6 (12.2%) | 5 (11.9%) | 11 (16.2%) |

\* The number of genes with a depth >1 is given in brackets. \*\* The number of genes with a mean base depth per 5 million reads (BD5M) >1 is given in brackets.

Combining all methods, we detected ARGs conferring resistance to 13 different classes of antibiotics, with two additional classes not detected with the SMS-based in-house analysis: Mupirocin and Quaternary Ammonium Compounds (QAC). The number of ARGs identified from the four individuals was 29 to 54 using the Ion AmpliSeq™ AMR panel, 14 to 30 using the Ion AmpliSeq™ Pan-Bacterial panel, and 9 to 28 using the SMS-based in-house analysis (Table 5). Two to seven ARGs were found with all three methods, while four to 16 ARGs were found with both the SMS-based in-house analysis and the AMR panel (Table S3). The AMR panel was the most sensitive, identifying 21 to 38 ARGs not found with any other method, representing 50-66.7% of the overall identified genes. Only two to three ARGs were identified with the Pan-Bacterial panel alone (4.4-7.1% of all genes) while 5 to 11 genes were identified with the SMS-based in-house analysis alone (11.9-16.2% of all the genes).

General performances of two methods (the SMS-based in-house analysis and the AMR panel) were compared in Figure 4.

**Figure 4.**
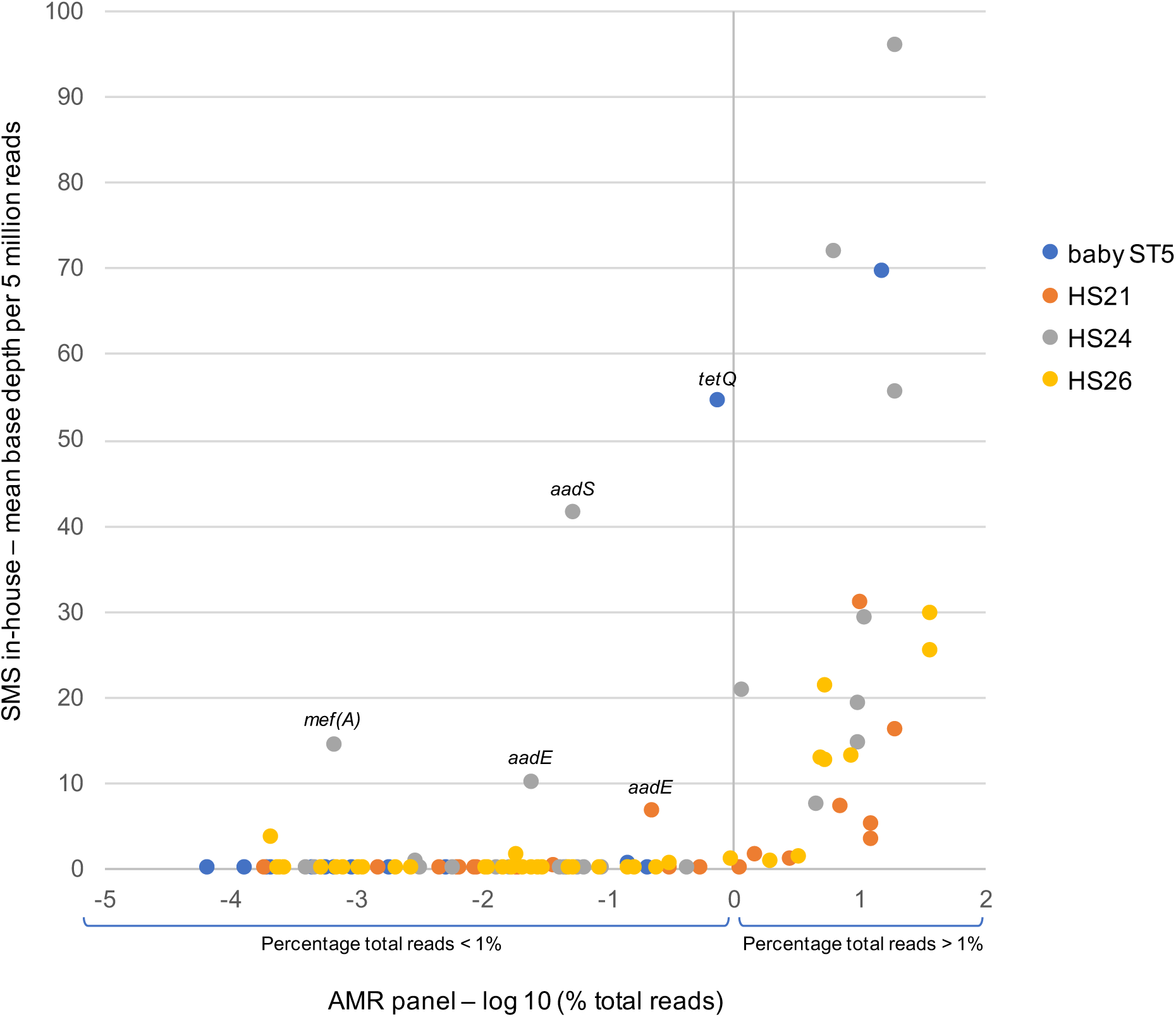
Performance summary of the in-house analysis from the shotgun metagenomics sequencing (SMS), compared to one targeted sequencing method using the Ion AmpliSeq™ AMR panel (Life Technologies). Results obtained with the AMR panel were analyzed with the One Codex online platform (available at: https://app.onecodex.com/); depth is defined as the average number of reads piled up across the entire gene.

## Discussion

Although only including a relatively low number of samples (n = 13), we identified a high number of ARGs (34 to 64) predicted to confer resistance to 13 classes of antibiotics. All sequencing methods were able to detect the most abundant genes, showing that ARGs were common in the gut microbiota of humans and dogs in Australia. Australian birds also carried ARGs in their gut, at a very low level in six wild Pacific black ducks from a non-urban environment, and at higher abundances in a Muscovy duck from an environment close to humans (semi-urban).

### ARGs conferring resistance to tetracycline, β-lactams and MLS_B_ are widespread

The most abundant ARGs present in the samples conferred resistance to tetracycline, β-lactams and MLS_B_ antibiotics. Interestingly, antibiotics within these classes are used frequently in human and veterinary medicine in Australia. Between 2015 and 2017, in human medicine, the most commonly dispensed antibiotics in community-based outpatient practice were β-lactams (57.3%), tetracycline (8.0%) and macrolides (5.1%), similar to those prescribed in hospitals (63.3% β-lactams; 8.2% tetracycline; 4.0% macrolides)^40^. Antimicrobial use in veterinary medicine in Australia is not as well documented, however the same three classes of antibiotics made up the majority (50.5 to 65.4%) of antimicrobials sold for animal use between 2005 and 2010 but ranked differently: macrolides and streptogramins (18.8 to 27.1%), followed by tetracyclines (16.3 to 28.6%) and β-lactams (9.2 to 14.6%)^41^. Studies have reported that subjects from countries with tighter policies on antibiotic usage in humans and animals have considerably less ARG levels^14^. In line with this, we would argue that the common usage of specific classes of antibiotics as noted above is likely driving the high abundance of matching ARGs in the gut microbiota of people and animals in Australia.

A large proportion of ARGs in our human samples were likely carried by bacteria within the Order Bacteroidales that is common in the human lower gut^42^. ARGs conferring resistance to tetracycline are one of the most commonly found in human gut microbiota^15^ and their prevalence has steadily increased in Bacteroidales since the 1970’s^43^. A study suggested that once acquired, ARGs to tetracycline can persist in Bacteroidales in the absence of antibiotic selection^43^. This could explain why ARGs to tetracycline were the most abundant in our data (detected in 12 samples out of 13, with up to 11 tetracycline ARGs in a single sample), despite tetracycline only being the third most common class of antibiotics prescribed in the Australian community behind several β-lactams^40^.

Mosaic tetracycline resistance genes, presumably arising by recombination between wild-type genes, have been discovered recently, and studies have shown that mosaic genes comprising tet(O), tet(W), and tet(32) sequences were abundant in DNA extracted from pig and human fecal samples^44^. In our data, mosaic genes tet (O/W), tet(O/32/O) and tet(W/32/O) were identified using the ResFinder tool. There is probably more undetected, as reads mapping against tet genes sometimes showed incomplete coverage with clear truncations. Targeted sequencing techniques might be required in order to properly assess tetracycline resistance that might otherwise be underestimated.

### Detected ARGs in humans and animals are diverse

The diversity of ARGs detected in humans was generally higher in adults compared to infants. This might be correlated with higher diversity of the gut microbiota in adults (the range of microorganisms found in the intestine changes dramatically between birth and the age of 3 years^45^) and/or a growing antibiotic selection pressure with age. Although the ARGs in infants were less diverse, they were at very high relative abundances, having the 2^nd^, 3^rd^ and 4^th^ highest mean coverage of ARGs (BD5M) of all the human samples. In Australia, antibiotic usage in children aged 0 to 9 years is almost equivalent to one antibiotic prescription per child and per year^46^, which is relatively high when compared to similar countries^47^. Although, the youngest infant still had ARGs evident when it had never been exposed to antibiotics.

The ARGs detected in the dog sample were somewhat similar to the ones detected in adult humans. This might reflect similarities in the diversity and composition of the dog and human fecal microbiota, with microbial exchange facilitated by direct contact^45^ as well as a similarity in the antibiotics used (as previously mentioned). Very few ARGs were detected in the Pacific black duck MAD (a pool of faeces from wild ducks collected in rural areas) compared to the Muscovy duck MUD (common duck collected in a semi-urban area). This is consistent with previous studies demonstrating that proximity to human activity increases the number of the antibiotic-resistant bacteria that are associated with wild birds. For example, gulls and geese nesting near waste or agricultural water harbor more antibiotic-resistant *E. coli* than do birds associated with unpolluted water^48,49^. Interestingly, ARGs to macrolide and tetracycline were the most abundant in the Muscovy duck MUD sample, overlapping with the antimicrobials used for therapeutic purposes in farm animals in Australia^41^. Most of the macrolides and tetracyclines are administered in food or water to pigs and poultry in Australian farms ^41^, and environmental contamination near farms with both antibiotics and ARGs has been demonstrated, as well as the higher abundance of ARGs in wild animals residing in the vicinity of farms^50^.

### The different sequencing methods have advantages and limitations

We tested three sequencing techniques and analysed the SMS-based data with three methods. The SMS-based in-house method and the AMR panel detected the greatest range of ARGs, with the highest diversity of ARGs obtained with a combination of the two methods. However, each method had advantages and disadvantages. The SMS-based in-house analysis and the AMR panel both detected abundant ARGs, while discordant results occurred for ARGs at low abundances, because of different detection sensitivities of the methods.

Discordant results between methods also occurred from the use of different AMR reference databases. Of the ARGs identified with the SMS-based in-house analysis only, many were absent from the AMR panel, and therefore could not have been detected by this method. Similarly, many of the ARGs identified with the AMR panel only were absent from the NCBI’s AMR database, therefore could not have been detected using the in-house analysis (Figure 4).

***The two targeted sequencing methods relying on AmpliSeq™ panels*** were very sensitive because of the pre-amplification step by PCR. The AMR panel identified the highest number of ARGs and captured >50% more genes not identified by the other methods. This included genes present in the NCBI’s AMR database but nonetheless undetected with the in-house method, e.g. erm(G), sat4, tet(M), lnu(C). Despite amplification, those genes were found with the AMR panel at a depth below 6,000 i.e. ≤ 1% of the total amplified reads per sample (see Table S3 for details). As a comparison, in sample HS24, all ARGs represented 0.2% of total SMS reads. With a non-targeted method, sensitivity can only be increased by increasing the number of reads.

On the contrary and less intuitively, four genes were detected at relatively high abundances with the SMS-based in-house method but at a low depth with the AMR panel: mef(A), aadE, aadS, tetQ (Figure 4). This might indicate poor efficiency of some primers. In sample HS24, the “One Codex” online platform reported a mean depth of 328 for tet(M) gene when in reality two sets of primers were amplified and detected at different depths (38 and 608). Other sets of primers may be working poorly, leading to false negative results or underestimated depths. Also, genes used to design the primers might also have been being quite different from the one present in our samples. This seemed to be the case for gene aadE detected in sample HS24, with the best match using the SMS-based in-house method (NG_047378.1 *Pediococcus acidilactici* aadE gene for aminoglycoside 6-adenylyltransferase) being only 69% similar to the aadE gene reference AF516335 used to design the AMR panel.

Targeted methods are limited by their inability to identify novel AMR determinants, whereas a functional SMS approach might provide more detail^52^. Using an in-solution probe-and-capture strategy^53^, researchers suggested that designed probes should target sequences with up to 15% nucleotide sequence divergence from a reference sequence, which would widen their applicability and target capacity toward newly characterized members of AMR gene families, which often differ from other members by only a few nucleotides^28^.

Because the depth analysis follows an amplification step, targeted methods are not fully quantitative compared to SMS. Nevertheless, it can be considered semi-quantitative as the depth generally gives a good idea of the initial quantity of the ARGs present.

***The SMS-based in-house analysis*** was not as sensitive as the targeted methods, being that the sequencing is random with no amplification step. However, (i) it is quantitative and (ii) it potentially allows identification of the bacteria carrying ARGs. Limitations of the SMS-based in-house analysis include reliance on manual checking of the ARGs detected. In the NCBI database used for the current study, references were constructed to include the ARG but also flanking regions of approximately 100 bp each side (if such regions were available in the original sequence report). This could trigger identification of false-positives for the presence of ARGs. That was the case for nimJ gene in HS24 that was originally identified at a BD5M = 0.3, but after manual checking, all reads mapped against the first 100 bp only of reference NG_048017 (*Bacteroides fragilis* nimJ gene for nitroimidazole resistance), i.e. the bacterial host genome.

Manual checking was also essential to avoid overestimation of a gene’s abundance. Gene tet(C) was identified in ST4-3mo at a BD5M = 13, but all reads mapped against a limited region of the gene and the consensus sequence was only 80.8% similar to the best reference in the NCBI’s AMR database NG_048174 (*Francisella tularensis* tet(C) gene for tetracycline efflux MFS transporter). The best match overall using BLASTN was gene tet(A) (98.9% similarity; reference MG904997). A tet(A) gene (reference NG_048158) was actually identified in ST4-3mo at a BD5M = 279.8, confirming that the reads mapped to NG_048174 / tet(C) were in fact not the best match.

Library post-amplification (done when the initial library quantity was too low) triggered unexpected problems and should be avoided when possible. Reference NG_047324 (*Escherichia coli* aadA1 gene for ant(3”)-Ia family aminoglycoside nucleotidyltransferase) was identified in MAD at a BD5M = 9 but the mapping result showed that all reads matched the same 200 bp region in the middle of the gene, so that the aadA1 gene coverage was only 22.7%. This specific sample (as well as ST4-3mo, MUD and DFS) was post-amplified, meaning that there might have only been a small amount of aadA1 gene in the initial DNA extract, and one fragment only was post-amplified and sequenced, leading to overestimating the depth of this gene in MAD.

In contrast, the depth may have been underestimated for some genes. In HS24, reference NG_047625 (*Bacteroides fragilis* cepA gene for beta-lactamase) was identified at a BD5M = 0.9 using a Q-value = 20, with NG_047625 being only 80.9% similar to the consensus sequence. This could be verified at a Q-value = 0, with good mapped reads, only quite different from the reference used for mapping; BD5M increased to 4.5, which is probably more accurate. One way to overcome this issue and obtain the real depth is to use the consensus sequence as the reference for re-mapping the reads. Identification from reads or contigs relies heavily on the reference databases, and this example shows that the closest match with a non-perfect reference can still allow the detection of a hypothetical ARG. But because only distantly related to the original curated ARG, further explorations are required to assess if the gene is actually a functional resistance gene.

Genes for which there is no good match in the reference database or in the NCBI general nucleotide database might be totally missed. When conducting read mapping coverage analyses at a Q-value = 0 instead of 20, we detected in some samples mixed populations of reads, i.e. one population matching the reference quite well, and the second population being more dissimilar. A coverage analysis at a Q-value = 20, showed the real depth of population 1, but discarded population 2 for which no better reference was available.

***The ResFinder tool***, especially when combined with the ***KmerResistance tool***, had a sensitivity similar to the SMS-based in-house analysis, albeit with the ARGs detected being slightly different. Even though the ResFinder and KmerResistance tools relied on the same database, results did not fully overlap, probably because they used different input data, i.e. contigs versus reads respectively. Compared to the in-house analysis, the online tools are more user-friendly and provide quick and relatively accurate results. Provided that the reference database is regularly updated to include missing genes (in our case the obvious ones being aminoglycoside/aadS, lincosamide/lnu(AN2), macrolide/mef(En2) and β-lactam/blaEC) these are valuable tools for scientists and diagnosticians requiring fast results. However, they don’t allow manually rechecking the mapping quality, and do not provide information on whether the genes are chromosomal or on plasmids, nor the taxa of the host bacteria. Even though the ResFinder tool provides an accession number corresponding to a good match for the host bacteria (from the contig sequence), it does provide a single match even though several other (potentially very different) species might show the same percentage of identity with the contig. Also, because we used the ResFinder tool with the contigs only, abundance is not provided, while the KmerResistance tool provides a depth result for the ARGs identified. The SMS-based in-house analysis provided the best accuracy regarding the abundance of ARGs, our calculated “BD5M” coverage taking into account the size of the gene as well as the depth of coverage.

***All the methods tested*** shared some common limitations. First, they rely on reference databases that are non-exhaustive, precluding the detection of ARGs very different to those included in the database, and impacting the abundance estimations of distantly related genes or when multiple similar genes are present. ARGs that show many different alleles were challenging to detect (e.g. cfxA / cfxA6 in sample DFS or aadA1 / aadA24 in sample MUD). For targeted methods, new (larger) panels are constantly being released, e.g. 37,826 probes targeting over 2,000 nucleotide sequences associated with AMR using a probe-and-capture strategy^28^, or 78,600 non-redundant genes (including 47,806 putative ARGs) using targeted metagenomics^51^.

Second, genotypic resistance does not necessarily reflect a phenotypic resistance. More studies comparing molecular data with phenotypic resistance are needed to assess how much of the detected resistome is linked to a functional resistant gene. It will also be important in the future to assess if those genes persist over time in an individual, and how much ARGs in the gut microbiota can be horizontally transferred to other bacteria species, potentially pathogenic ones.

This study is the first to describe the bacterial resistome in humans and animals from Australia, comparing and assessing various methods. A high diversity of ARGs was identified, as well as a high variability between samples, even with a limited number of samples analysed. We show that resistomes from (human) adults and infants are dissimilar, and that resistomes from adults, dogs and urban birds are more similar than that from rural birds. Ongoing surveillance of ARGs abundance in people and animals (including livestock and wild animals) in Australia using the techniques described here could help inform policy makers and health care professionals on the most prudent use of these important drugs to ensure we have access to effective treatments of bacterial infections well into the future.

### Ethics statement

Ethics approval for the use of seven pregnant mothers samples was granted by the Barwon Health Human Research and Ethics Committee (HREC 10/24). The study involving the three infant samples was deemed negligible or low risk by the Barwon Health Human Research Ethics Committee and therefore exempt from full committee review (HREC approval 17/119). Bird sample collection was approved under Deakin University’s Animal Ethics Committee project number B43– 2016 and Department of Environment, Land, Water and Planning permit number 1008206. Fecal samples from the dogs were environmental samples collected on the ground and did not require ethics approval.

## Data availability

All data generated or analysed during this study are included in the published article.

## Acknowledgements

This work was supported by funding from Deakin University, Barwon Health and CSIRO and from NHMRC Equipment Grant GNT9000413 and from the NHMRC Australian Partnership for Preparedness Research on Infectious Disease Emergencies (APPRISE) CRE to SA. We thank the Barwon Infant Study (BIS) Investigator group: Peter Vuillermin, Anne-Louise Ponsonby, Mimi LK Tang, Richard Saffery, David Burgner, John Carlin, Sarath Ranganathan, Katie Allen, Peter Sly, Len Harrison and Terence Dwyer for providing access to fecal samples from humans and we are grateful to Prof Marcel Klaassen for providing the duck samples, to Jessie Vibin for DNA extraction of the two duck samples, and to Aseel Norladin for help in processing the samples during the libraries preparation.

## Author Contributions

Project design and coordination was performed by SA. Samples were collected by and selected by SA and FC, with adult human samples collection coordinated by the BIS investigator group. Nucleic acid was extracted by VG and FC. VG and FC performed the sample library and NGS DNA sequencing. VG, AC and SA performed the reads analyses. VG wrote the initial manuscript with input from SA, and later versions were based on input and suggestions from all.

## Legends to Supplementary Tables

**S1 Table: Detailed results from the mapping of Shotgun Metagenomic Sequencing (SMS) against the NCBI’s AMR reference gene database using the Torrent Suite TMAP plugin.** Reads were queried against the database using Magic-BLAST and contigs were queried against the database using BLASTN. For each reference (line), red squares indicate the samples for which the reference was selected from the reads and/or contigs as the best (or one of the best), while the other samples were mapped against the reference just as a comparison with the other samples. Mean base depth per 5 million reads (BD5M) is highlighted in yellow when between 1 and 10, or red when > 10. When a reference is the best mapping result but the number of reads mapped to the reference is low (< 20), the exact number of reads is provided in brackets.

**S2 Table: Detailed comparison of the analyses with the SMS-based method using (i) the KmerResistance tool, (ii) the ResFinder tool and (iii) our in-house method.** For ResFinder and KmerResistance the percentages provided are the template coverage. For the in-house method, the calculated “mean base depth per 5 million reads” is provided. * gene not in the NCBI’s AMR gene database; § gene not in the ResFinder database.

**S3 Table: Results from a SMS-based in-house method and a targeted metagenomic sequencing method using the Ion AmpliSeq™ AMR panel (Life Technologies).** Results obtained with the AMR panel were analysed with the One Codex online platform (available at: https://app.onecodex.com/); depth is defined as the average number of reads piled up across the entire gene. SMS: shotgun metagenomics sequencing; BD5M: mean base depth per 5 million reads; MLS: macrolide, lincosamide, streptogramin B; QAC: quaternary ammonium compounds; gni: gene not included in Bioproject database; gnp: gene not included in the AMR panel. Genes detected with both methods are highlighted in green; genes in blue are detected with the AMR panel only; genes in yellow are detected with the SMS method only.

